# Accurate and interpretable gene expression imputation on scRNA-seq data using IGSimpute

**DOI:** 10.1101/2023.01.22.525114

**Authors:** Ke Xu, ChinWang Cheong, Werner Pieter Veldsman, Aiping Lyu, William K. Cheung, Lu Zhang

## Abstract

Single-cell RNA-sequencing (scRNA-seq) enables the quantification of gene expression at the transcriptomic level with single-cell resolution, enhancing our understanding of cellular heterogeneity. However, the excessive missing values present in scRNA-seq data (termed dropout events) hinder downstream analysis. While numerous imputation methods have been proposed to recover scRNA-seq data, high imputation performance often comes with low or no interpretability. Here, we present IGSimpute, an accurate and interpretable imputation method for recovering missing values in scRNA-seq data with an interpretable instance-wise gene selection layer. IGSimpute outperforms ten other state-of-the-art imputation methods on nine tissues of the Tabula Muris atlas with the lowest mean squared error as the chosen benchmark metric. We demonstrate that IGSimpute can give unbiased estimates of the missing values compared to other methods, regardless of whether the average gene expression values are small or large. Clustering results of imputed profiles show that IGSimpute offers statistically significant improvement over other imputation methods. By taking the heart-and-aorta and the limb muscle tissues as examples, we show that IGSimpute can also denoise gene expression profiles by removing outlier entries with unexpected high expression values via the instance-wise gene selection layer. We also show that genes selected by the instance-wise gene selection layer could indicate the age of B cells from bladder fat tissue of the Tabula Muris Senis atlas. IGSimpute has linear time-complexity with respect to cell number, and thus applicable to large datasets.

## Introduction

The development of single-cell RNA sequencing (scRNA-seq) technologies has made it feasible to investigate cellular heterogeneity at the single-cell resolution. Compared to bulk RNA-seq, scRNA-seq measures gene expression profiles of individual cells instead of average values of a group of cells. It has been applied to identify novel cell types and their lineage development [1–5]. Furthermore, scRNA-seq can help better deconvolve and interpret tissues with high cellular heterogeneity, such as tumor microenvironments, providing a practical way to elucidate the interplay between genes, cells and diseases [6–8]. The mainstream scRNA-seq platforms are either droplet-based [9–11] or plate-based [12–14]. The two approaches have trade-offs between sensitivity and throughput [15]. The plate-based platforms capture low abundance RNA molecules for each cell with high sensitivity, but can only be applied to a small proportion of cells due to cost constraints. Droplet-based platforms are popular because of their lower costs per cell, enabling exploration of a large number of cells simultaneously [15]. However, the analysis of droplet-based scRNA-seq is challenged by artificial zeros (dropouts [16]), which are difficult to be distinguished from the biological zeros (the latter arises from RNA molecules with extremely low abundance). For example, the dropout rates of two housekeeping genes *ACTB* and *GAPDH* in all tissues sequenced by droplet-based scRNA-seq from the Tabula Muris atlas are approximately 13.8% and 31.0%, respectively. These unexpected zeros substantially affect the performance of downstream analysis (**Supplementary Figure 1** and **Supplementary Note 1**).

Many methods have been developed to recover dropouts by considering gene-gene and cell-cell similarities. KNN-smoothing [17] applies a variance-stabilizing transformation to calculate cell-cell similarities by assuming scRNA-seq data follow a Poisson distribution. MAGIC [18] transforms a cellcell distance matrix to a normalized Markov matrix with Gaussian kernels and applies random walk to aggregate information from neighboring cells. SAVER [19] estimates the parameters of a Poisson-Gamma mixture distribution using non-zero entries in gene expression matrices and imputes dropouts based on this distribution. ScImpute [20] estimates the cell-specific gene dropout probabilities based on a Gamma-Normal mixture distribution generated by spectral clustering. WEDGE [21] imputes dropouts using a biased low-rank matrix decomposition to reconstruct cell-wise and gene-wise correlations. Recently, some tools have adopted advanced deep learning techniques to implicitly consider gene-gene and cell-cell similarities. Deep count autoencoder (DCA) [22] introduces an autoencoder with a zero-inflated negative binomial (ZINB) distribution to impute scRNA-seq data. DeepImpute [23] accelerates imputation by splitting genes into groups and imputing dropouts for each group parallelly using their highly correlated genes. SAUCIE [24] is a multitask framework that performs dropout imputation, batch effect removal and cell-type clustering by introducing novel loss functions in the autoencoder, including reconstruction loss, maximal mean discrepancy loss and information dimension loss. ScGNN [25] applies graph neural networks with a left-truncated Gaussian mixture model to model cell-cell similarities and heterogeneous gene expression profiles iteratively. scVI [26] develops a variational autoencoder by assuming ZINB distribution in the latent space and decouples the library size (i.e. the expression summation of all genes in a cell) to distinguish artificial and biological zeros.

Off-the-shelf imputation methods commonly adopt global feature selection [27–30] (i.e, selecting the same subset of genes across all cells) as an independent data pre-processing step to select highly variable genes or to remove the genes expressed in just a few cells. These methods calculate the gene-gene and cell-cell similarities using all entries in gene expression matrix, and all entries are considered equally important. However, we found some local outlier entries in the gene expression matrix (e.g., some cell type marker genes expressed in cells that are not from the corresponding cell type) could be hidden among highly variable genes (**Supplementary Figure 2** and **Supplementary Note 1**). Global feature selection is unable to remove such noise. Instance-wise feature selection [31, 32], in contrast, allows selection of specific genes in cells and has the potential to reduce the impact of local noise.

In this paper, we accordingly present IGSimpute as an accurate and interpretable imputation method with a novel application of instance-wise gene selection for imputing missing scRNA-seq data. IGSimpute includes a denoising autoencoder with an instance-wise gene selection layer as well as a gene-gene interaction layer to allow more robust estimation of cell-cell and gene-gene similarities (**Figure 1**). For each cell, the instance-wise gene selection layer selects the genes that are to contribute to the construction of cell embedding and the estimation of gene-gene similarities. We compared IGSimpute with ten other state-of-the-art imputation methods using 12 scRNA-seq datasets. IGSimpute achieved the top rank on nine of the 12 datasets and was ranked among the top three methods on the other three datasets. We show that usage of the gene expression matrix imputed by IGSimpute significantly improves downstream analysis such as cell type clustering and visualization. We also show that the genes selected by the instance-wise gene selection layer are candidate markers of biological characteristics such as age.

**Figure 1.**
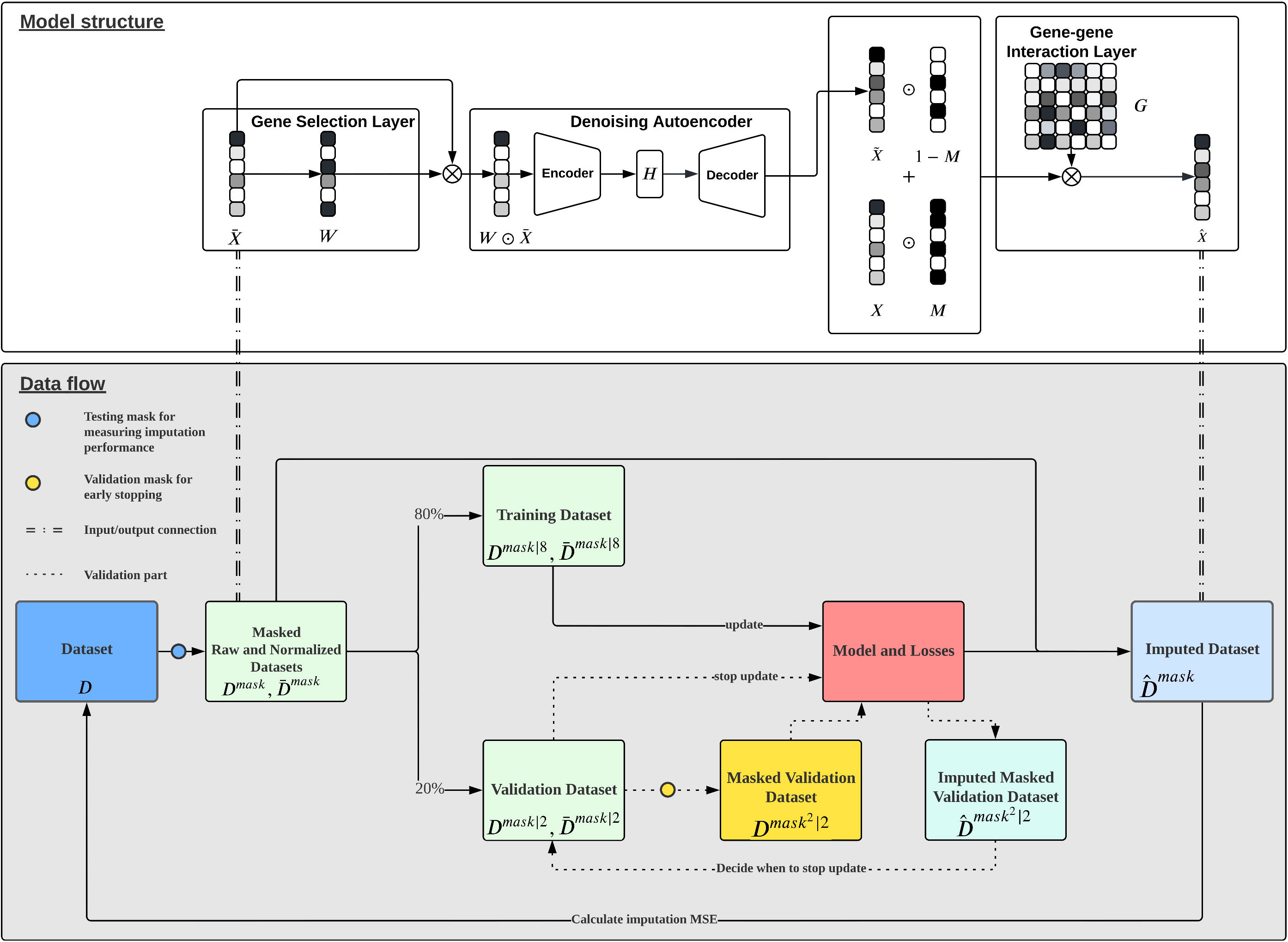
IGSimpute model structure and data flow diagram. (**Top panel**) The normalized input matrix 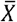 is first used to obtain a weight matrix *W*. The Hadamard product of both matrices is then taken as the input of a denoising autoencoder for generating the imputed expression values 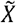. We then combine two matrices by keeping nonzero expression values of the raw input matrix *X* and replacing the others entries (i.e. zero entries) with the imputed expression values from 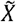. The composed matrix is finally multiplied by the gene-gene interaction layer to get the final imputed expression matrix 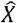. (**Bottom panel**) The raw dataset is first masked such that we can assess the imputation performance for the masked entries. We further normalize the masked raw dataset and split the masked datasets into the training dataset and validation dataset. The validation dataset will be masked again but by the model such that model can assess the imputation performance internally and decide when to stop training. The imputed dataset is then compared to the raw dataset to calculate imputation MSEs.

## Material and methods

### Sourcing of scRNA-seq datasets

We downloaded the scRNA-seq derived gene expression count matrices of 12 tissue datasets (**Supplementary Table 1**) in the Tabula Muris atlas [33] in order to compare IGSimpute with the other imputation tools. To investigate the interpretability of the instance-wise gene selection layer, we also downloaded the scRNA-seq derived gene expression count matrix of a brown adipose tissue (BAT) dataset in the Tabula Muris Senis atlas [34]. More details on the sourced datasets are available in **Supplementary Note 2**.

### Pre-processing of scRNA-seq data

We used Scanpy [27] to perform pre-processing and quality filtering on the gene expression count matrix of each tissue dataset. For the Tabula Muris atlas, we first extracted the cells from each tissue dataset using associated metadata and excluded genes/cells from further analysis if (i) genes were expressed in less than three cells (using scanpy.pp.filter_genes with min_cells = 3); (ii) cells with 500 or less expressed genes (using scanpy.pp.filter_cells with min_genes = 500); and (iii) cells with insufficient library size defined as less than 1,000 expression counts (using scanpy.pp.filter_cells with min_count = 1000). For the BAT dataset, we extracted B cells and removed (i) External RNA Controls Consortium (ERCC) [35] spike-ins; and (ii) genes with expression counts less than ten (using scanpy.pp.filter_genes with min_counts = 10).

We normalized cell library sizes (scanpy. pp.normalize_per_cell) to make the total gene expression counts comparable and transformed gene expression values logarithmically to mitigate extreme values in the count matrices (using scanpy.pp.log1p). Highly variable genes were identified using scanpy.pp.highly_variable_genes with the “seurat” [28] approach. Raw counts of the highly variable genes were then extracted from the original gene expression matrices to generate data matrix *D* for subsequent analysis. To evaluate the performance of imputation, we created a testing mask that randomly zeroed out 20% of the non-zero entries from *D*. We multiplied the testing mask by the data matrix *D* as *D^mask^*. To access the robustness of imputation, we replicated the procedure 10 times. All imputation methods take raw (*D^mask^*) gene expression matrices as input and IGSimpute further performs Z-score normalization to obtain normalized gene expression matrices 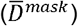, while other imputation methods will normalize *D^mask^* internally if they have relevant options since not all imputation methods can accept 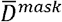 as input.

### The network structure of IGSimpute

IGSimpute implements an autoencoder with an instance-wise gene selection layer and gene-gene interaction layer to improve scRNA-seq imputation and downstream analysis (**Figure 1**). The instance-wise gene selection (IGS) layer is inserted between the input layer and the encoder to remove the non-informative or redundant genes that contribute little to cell embedding. The IGS layer generates a gene weight matrix *W* (*W* ∈ [0,1]), given as

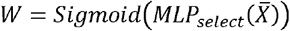

where MLP() is a multilayer perceptron and 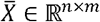 denotes the normalized gene expression matrix with n and m being the number of cells and genes respectively. The cell embedding is generated by the encoder as

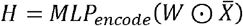

IGSimpute includes also a gene-gene interaction layer between the decoder and the final output layer which composes a score matrix *G* where the values in *G* indicate the strengths of gene-gene interactions. The score matrix *G* is learnable so that the expression counts of the genes can satisfy the following equation:

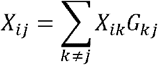

where *X* is the raw gene expression matrix. This essentially assumes that gene counts can be well estimated by the weighted sum of the counts of the interacting genes.

The imputed gene expression matrix 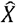 can thus be computed by

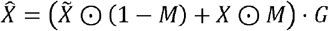

where

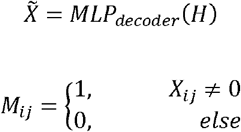

We inserted Gaussian noise and dropout layers between the input and instance-wise gene selection layer to increase model robustness. The detailed module structure and parameters are described in **Supplementary Note 3**.

The intuition behind the network structure is that more robust cell embedding can be generated by selecting fewer but more informative genes. By applying an instance-wise gene selection layer, the cell-cell similarities will be determined by gene subsets that are robust to random dropouts and outliers. A gene-gene interaction layer is set up to capture the global gene-gene similarities. The gene-gene interaction layer is data-driven and has no distribution assumption.

### Training Scheme

We obtained *D^mask^* to compare IGSimpute with the other imputation methods (**Figure 1**). IGSimpute distributes the cells of *D^mask^* into training and validation sets with the ratio of *D*^*mask*|8^:*D*^*mask*|2^=8:2. The loss functions involved in the training step are:

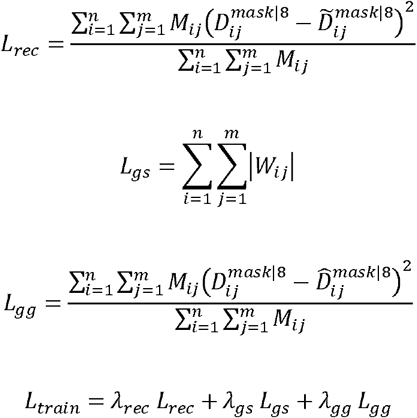

where *n* and *m* are the total numbers of cells and genes, respectively.

The reconstruction loss *L_rec_* encourages the output of the decoder to reconstruct *D*^*mask*|8^. The regularization loss *L_gs_* is applied to the instance-wise gene selection layer to make the gene weight matrix *W* sparse to avoid overfitting. Another reconstruction loss *L_gg_* is designed to minimize the imputation errors after gaining weights from the gene-gene interaction layer. We further generate a validation mask to zero out 20% of non-zero entries in *D*^*mask*|2^ as *D*^*mask*^2^|2^ to determine the number of training epochs for early stopping based on the model’s performance. The training procedure would be terminated before achieving the maximum allowed epochs if the MSE was not decreased after 1,000 epochs for the Tabula Muris datasets and 100 epochs for the mouse brown adipose tissue dataset. IGSimpute was implemented in Python 3 using Tensorflow [36] 1.15 and adopted AdamW [37] with a weight decay of 0.0001 and a learning rate of 0.0001 to accelerate model convergence. The batch size was set to 256 for all datasets.

In the training stage, we took 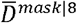 as the input for the neural network and optimized *L_train_* using *D*^*mask*|8^. During the validation stage, we only need to feed the input 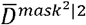 and obtained the output 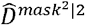 without optimizing any loss function. We calculated the MSE between *D*^*mask*|2^ and 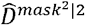 for the early stopping purpose as illustrated above. We used *D^mask^* as the input in the testing stage and extracted the output 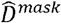 for post-processing.

### Post-processing

The negative values in 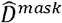 were converted to zeros to avoid unexpected values in the output gene expression matrix and the non-zero entries in *D^mask^* would not be imputed.

We used a KNN (K-Nearest Neighbors)-based post-processing strategy to recognize possible biological zeros and to remove corresponding imputed expression values if most of the neighbors had low expression values. The strategy consisted of three steps: (i) we extracted the expression profiles for each gene across all cells and labeled low expression values if they were zeros or included in the smallest 20% of non-zero entries; (ii) only the non-zero entries for a particular cell would be used to calculate the Euclidian distances with its neighbors (ten nearest neighbor candidates for each cell); (iii) we converted the imputed gene expression values to zeros if the same gene was labeled as lowly expressed in more than 80% of the neighboring cells.

### Imputation Comparison Settings

We provided *D^mask^* as the input for all methods except IGSimpute because some methods had built-in normalization steps (**Supplementary Note 4**). Each imputation method was run with the same group of parameters for all 12 datasets unless the imputation method (e.g., scGNN) required the corresponding cluster number in which case the ground truth cluster number would be given.

### Cell type clustering

To assess the influence of different imputation methods on downstream analysis, we performed cell type clustering on the imputed gene expression profiles using SC3 [38], where the number of clusters was from the ground truth. We converted the negative values in the imputed gene expression profiles to zeros followed by normalizing them logarithmically with base 2. We disabled the commands to filter out genes and cells in SC3 to obtain the clustering for all cells.

### Evaluation Metrics

For each scRNA-seq dataset, we repeatedly generated a randomly masked count matrix and imputed it in turn with ten different seeds to validate robustness. With each turn, the order of cells and genes in *D^mask^* was shuffled to reduce the chance of using similar mini-batches for model training, which would further increase the difference between replications.

We used mean squared error (MSE) and adjusted rand index (ARI) [39] to evaluate the performance of imputation and clustering, respectively. MSE was calculated by comparing the difference between the raw and imputed gene expression count matrix in the testing masked entries as follows:

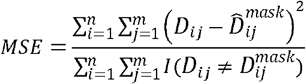

where *I* is the indicator function.

ARI measured how well the generated clustering matched with the benchmark ranging from −1 to 1.

Higher ARI indicates better clustering. Given the ground truth partition *A* = (*a*_1_, *a*_2_, …, *a_t_*) and the predicted clustering partition B = (*b*_1_, *b*_2_ …, *b_p_*), ARI could be defined as

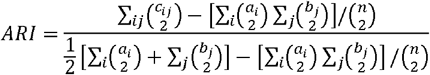

where 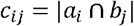 and n denoted the total cell number.

We calculated p-values for each dataset and combined them using scipy.stats.combine_pvalues using Fisher’s method. To compare the mean and variance of MSE, we used scipy.stats.wilcoxon (i.e. Wilcoxon signed rank test) with alternative=“less” and scipy.stats.kruskal (i.e., Kruskal-Wallis H-test) to generate p-values for ten replications of each dataset. To compare the mean ARI, we calculated p-values for the three best replications of each dataset using scipy.stats.ttest_ind with equal_var=False (i.e. Welch’s t-test).

### Heatmap visualization

We selected genes for heatmap visualization in the following steps. Firstly, we applied scanpy.pp.log1p on the raw count profile and used scanpy.tl.rank_gene_groups with method=‘t-test’ to identify the candidate characterized genes for cell types. We selected genes with log (fold change) ≥ 3, q-value≤ 0.05 and average expression ≥ 1 for each cell group and took the union of these gene lists as a candidate gene list. Secondly, we selected the top three genes for visualization from the candidate gene list for each cell type by sorting the multiplication of the weight matrix and the normalized gene expression. Finally, we plotted heatmaps using scanpy.pl. heatmap and we set vmin=-3 and vmax=3 for better visualization.

### B cell development visualization

We first performed the principal component analysis with seed 42 to reduce the dimensions of the normalized expression matrix to 50 and used UMAP with the same seed to further reduce the dimensions into two. We aggregated the weights in *W* for each gene across all cells and selected the top 30 genes with the largest weights as core genes. For each cell, the weights of these 30 genes were aggregated to calculate SUM(*W*).

## Results

### IGSimpute outperforms the other methods in imputing dropouts of scRNA-seq data

We compared the performance of IGSimpute to 10 other state-of-the-art imputation methods (**Methods**) using 12 tissue datasets from the Tabula Muris atlas. We generated a testing mask to treat 20% of entries of gene expression matrices as “artificial dropouts” to evaluate their imputation performance by calculating the divergences between imputed and true values (**Methods**). This procedure was repeated ten times to alleviate the influence of unstable results. We found IGSimpute achieved the lowest average MSE and variance in 9 out of 12 datasets, which were significantly better than the second-to-best methods (scVI with p-value= 1.58 × 10^−12^ for the MSE mean; DeepImpute with p-value= 2.16 × 10^−10^ for the MSE variance, **Methods**). Although IGSimpute was not the best in the other three datasets (heart-and-aorta, limb muscle and trachea), it still ranked in the top three and was comparable with the best results (average difference: 4.41%) (**Figure 2**). In addition, we found that only IGSimpute, SAUCIE and MAGIC were always better than baseline across all tissues while other imputation methods might produce worse imputation results than baseline in some tissues. For example, scVI had higher MSEs in thymus tissue dataset and other imputation methods had higher MSEs than baseline in at least two datasets (DCA in lung dataset and thymus dataset, DeepImpute in heart and aorta dataset, kidney dataset, spleen dataset and trachea dataset, kNN-smoothing in all datasets, scImpute in all datasets except marrow dataset, WEDGE in heart and aorta dataset, liver dataset and spleen dataset). We did not summarize SAVER and scGNN in the analysis above since they nearly did not impute any zero entries such that they had nearly identical bar height with baseline. SAVER and scGNN might prefer imputing low expression entries instead of zero expression entries. The imputation results demonstrated that IGSimpute was robust to data variations and could always give better imputation compared to baseline. We also used scatter plots to visualize the imputation results on bladder tissue and examined whether the imputed gene expression profiles were unbiased (**Figure 3A**). We observed that IGSimpute and DeepImpute could unbiasedly impute the gene expression values. scImpute and kNN-smoothing were observed to overestimate the masked gene expression values while the other six tools (DCA, SAUCIE, SAVER, scGNN, MAGIC and WEDGE) underestimated them.

**Figure 2.**
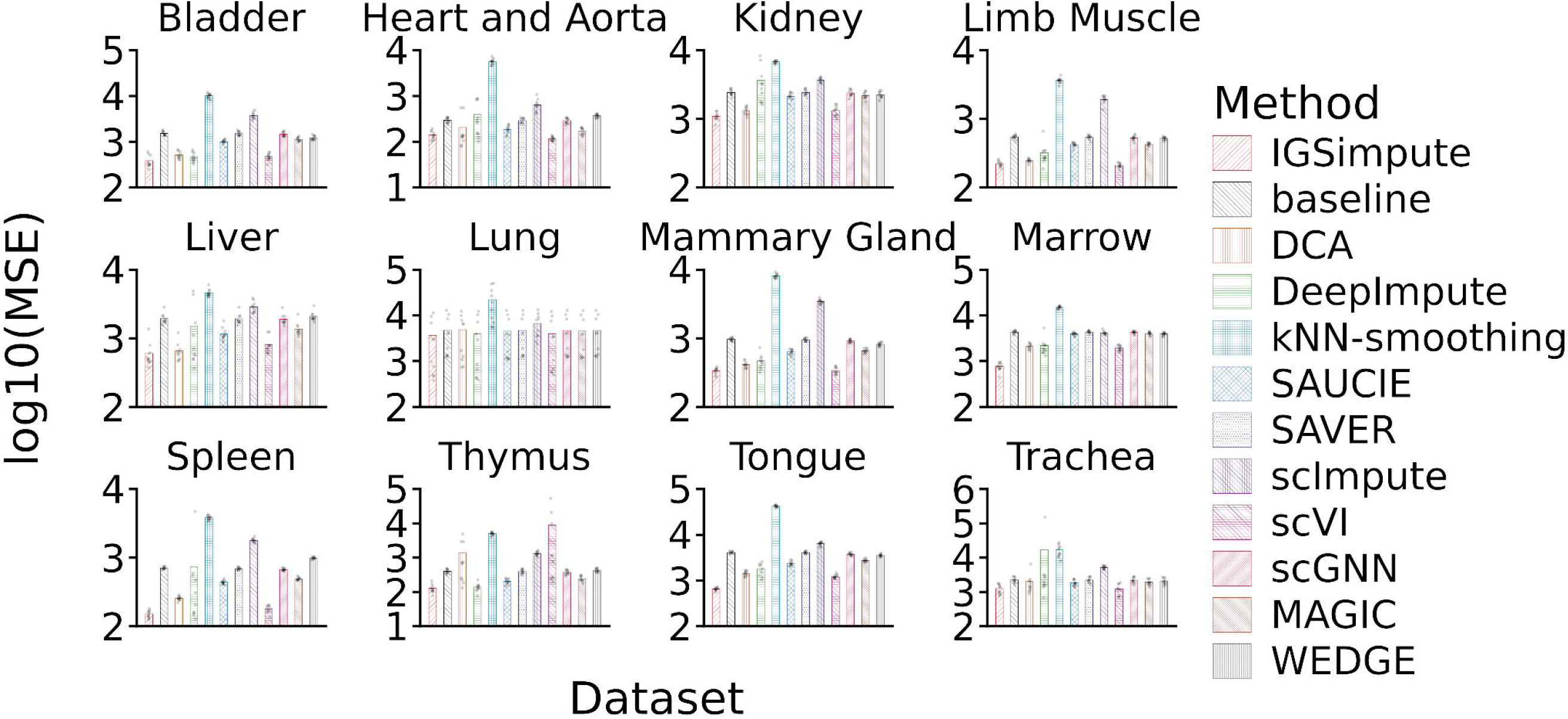
The imputation performance of IGSimpute and ten other state-of-the-art methods on twelve scRNA-seq datasets in the Tabula Muris atlas. The dots within the bars are the MSEs of the replicates and the bars represent the average of the MSEs across ten replicates for each method. Lower MSEs indicate better imputation performance.

**Figure 3.**
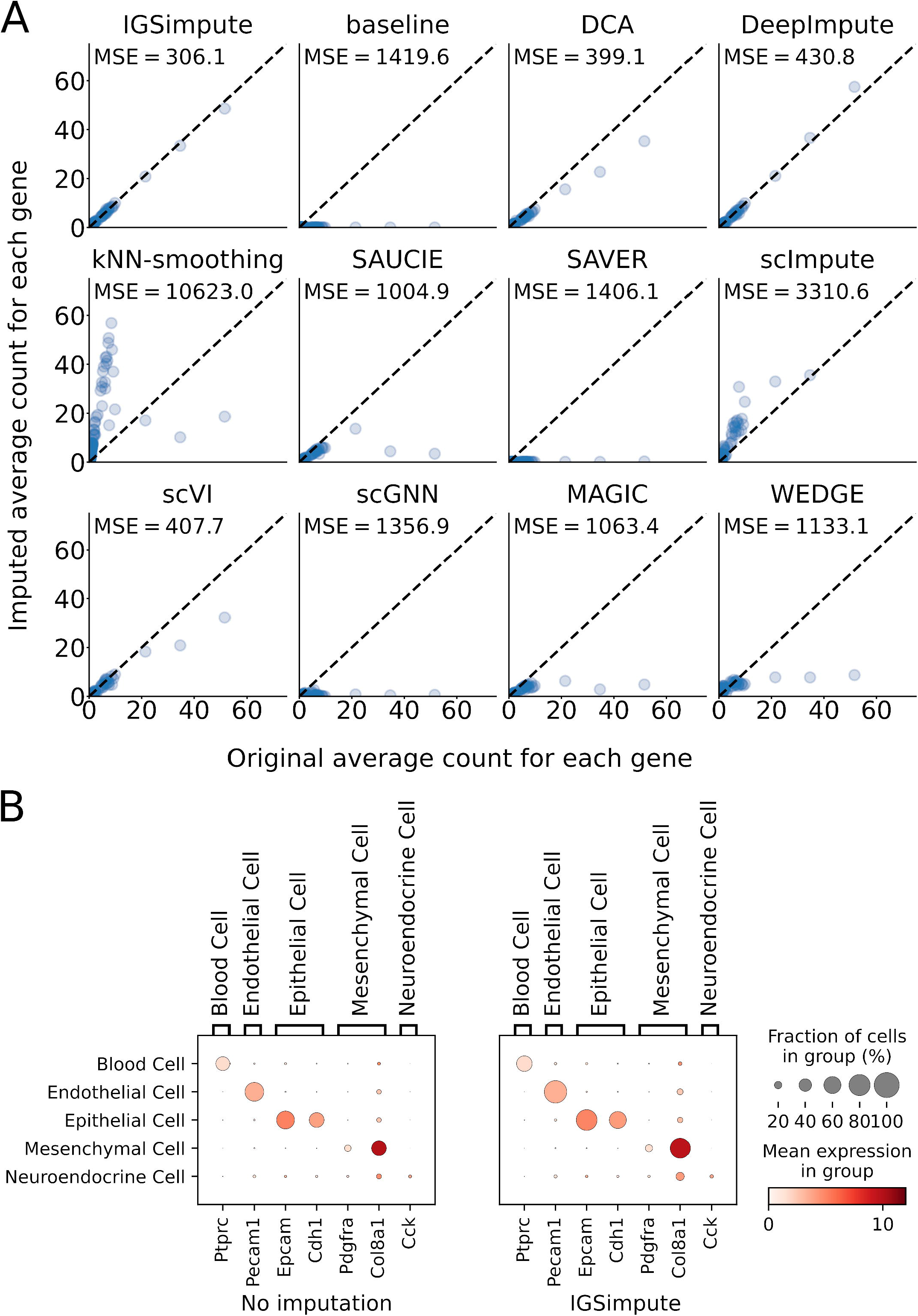
IGSimpute achieved unbiased imputation on bladder tissue and imputed the missing values of marker genes of trachea tissue with high accuracy. (**A**) Comparison of the difference between imputed and true gene expression values of bladder tissue data. Each point denotes a gene, where its horizontal and vertical coordinates represent the true and imputed gene expression values, respectively. The points above or below the diagonal lines suggest overestimating or underestimating corresponding gene expression values. (**B**) Gene expression profiles of the marker genes from five trachea tissue cell types before and after imputation. Circle size and color represent the percentage of non-zero expression entries and the mean expression values, respectively. The color mapping was truncated by 10 for better visualization.

Cell-type specific marker genes should have similar gene expression patterns in the corresponding cell types. We selected seven marker genes [33] from five cell types in trachea tissue to examine whether marker genes can be recovered (**Figure 3B**). We found the percentages of marker genes with non-zero expression values increased from 42.88% to 54.37% on average in their corresponding cell types. The imputed and original gene expression values of marker genes were quite close (average difference: 2.54%), suggesting IGSimpute achieved high confident imputation on the marker genes. For example, *Epcam* is a marker gene of epithelial cells and its expression was 5.06 and 5.21 before and after imputation and the percentage of dropouts reduced from 35.99% to 20.29%.

### IGSimpute improves cell type clustering

The aim of dropout imputation was not only to fill in the missing gene expression values but also to improve downstream analysis such as cell type clustering. We applied a scRNA-seq clustering method SC3 to compare cell type clustering before and after imputation (**Figure 4**), where K was assigned the same value as the ground truth in the benchmark for all the methods. We observed the gene expression imputed by IGSimpute achieved the highest adjusted rand index (ARI) for cell type clustering which is significantly better than using baseline data (without imputation, p-value= 1.6 × 10^−8^, **Methods**) and the second-to-best algorithm (with imputation, DeepImpute, p-value= 6.52 × 10^−3^, **Figure 4A** and **Supplementary Figure 3**). The cell embedding of trachea tissue was illustrated with the use of UMAP plots [40] to demonstrate the impact of imputation on cell type clustering. We observed that IGSimpute, DCA, DeepImpute, scImpute, scVI and WEDGE could improve the cell embedding and that the cells from different cell types were rarely mixed in the UMAP plots (**Figure 4B**). In addition, all imputation methods except IGSimpute failed to connect epithelial cells that spread over the whole embedding space together.

**Figure 4.**
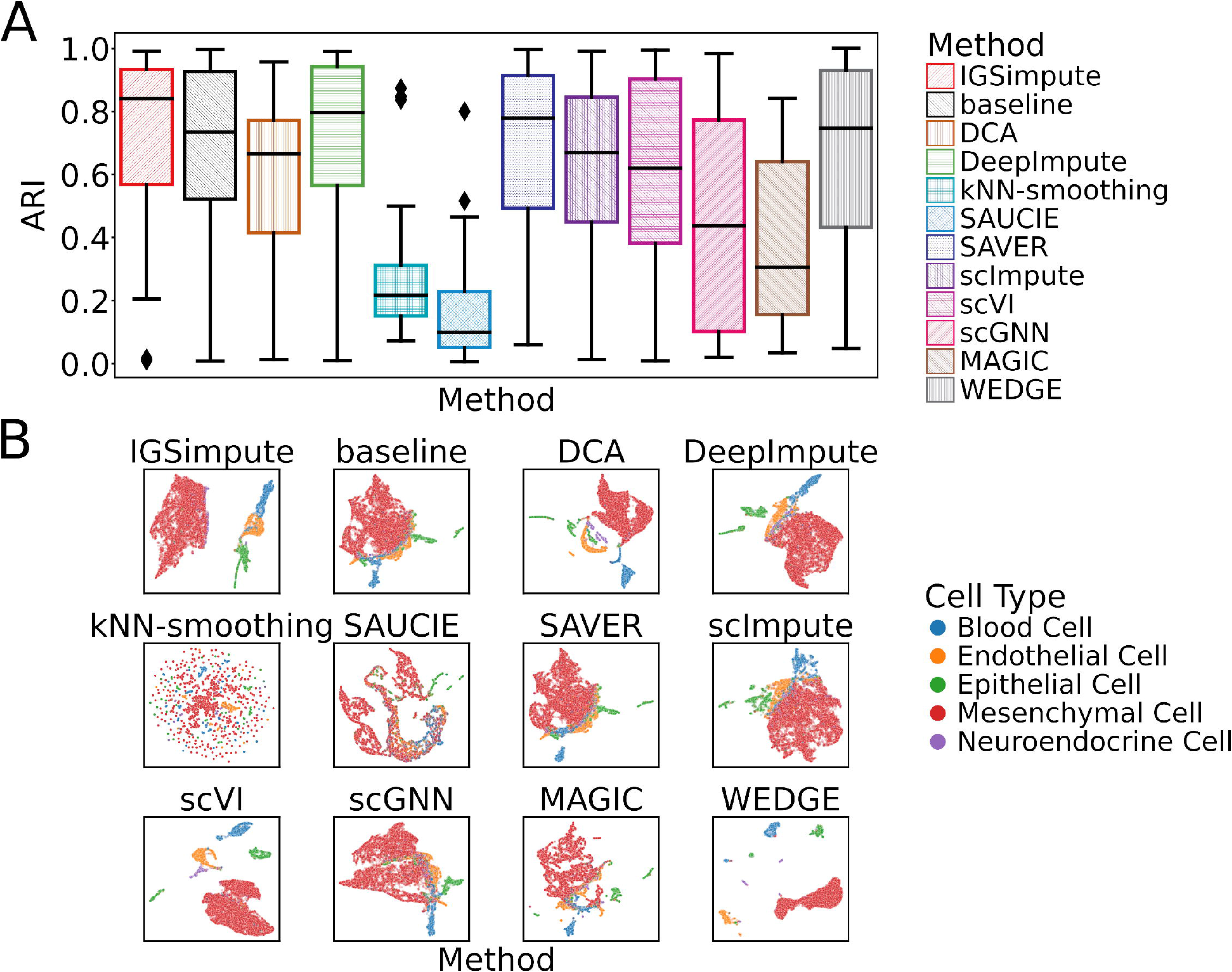
Clustering of the imputed profiles of IGSimpute achieved the highest median ARI on average across all 12 tissues in the Tabula Muris Atlas and visualization of the imputed profiles of IGSimpute showed better separation compared to other methods. (**A**) The box plots depict the clustering results for all 12 tissues. Boxes are color-coded with the respective methods. The horizontal line is the median value for each respective group of clustering results. For most of the methods and datasets, the same parameters were used when performing clustering by SC3. (**B**) UMAP plots of 12 imputed profiles from each imputation method for the trachea dataset. IGSimpute embedded cells comparatively well as indicated by the delineation of the clusters on its plot. ARI, adjusted rand index. UMAP, uniform manifold approximation and projection.

### Instance-wise gene selection denoises the gene expression matrix

The instance-wise gene selection layer is a key component of IGSimpute, which calculates the weights of genes in different cells to define their importance in achieving accurate imputation and reconstructing the gene expression matrix. We found that this layer could reduce the weights of less important genes in the cells, especially for the outlier highly expressed genes (**Figure 5A-D**).

**Figure 5.**
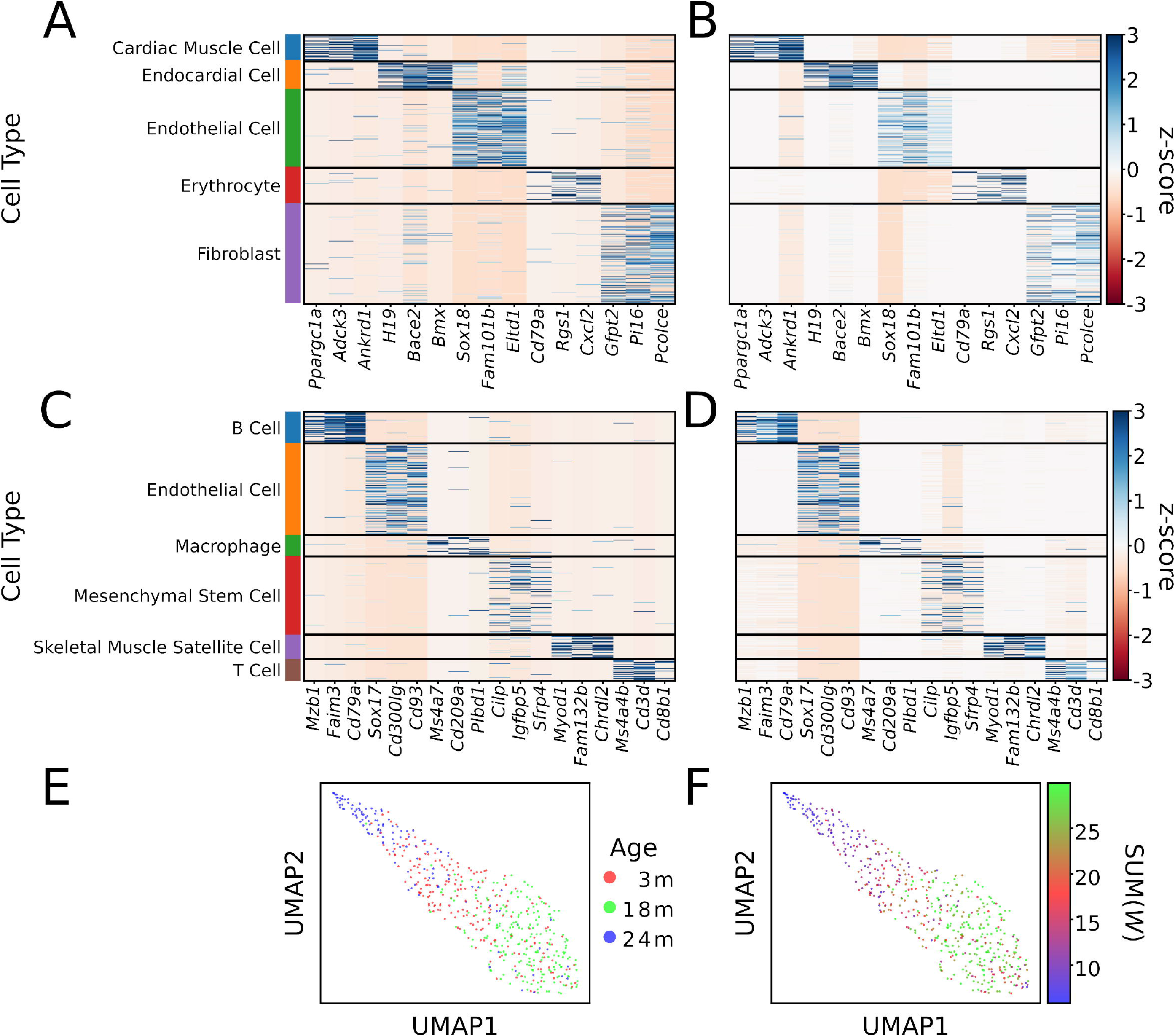
Instance-wise gene selection denoises scRNA-seq gene expression matrices and improves model interpretability. (**A-B**) Normalized and denoised gene expression matrices of the heart and aorta tissue and (**C-D**) the limb muscle tissue. Each column and row correspond to a gene or a cell, respectively. (**E**): UMAP plot for B cells with color indicating the mouse age. (**F**): UMAP plot with color indicating the values of SUM(W). The sum of aggregated core gene weights SUM(*W*) perfectly represents the B cells from mice aged 3, 18 and 24 months.

The denoised gene expression matrix could improve downstream analyses due to its better signal-to-noise ratio. We visualized the normalized 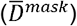 and denoised gene expression matrices 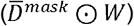 of the heart-and-aorta (**Figure 5A** and **B**) and limb muscle tissues (**Figure 5C** and **D**) to examine whether *W* (that is learned in the instance-wise gene selection layer) could help denoise outlier entries. We examined the top three characterized genes (**Methods**) of each cell type and found these genes were all highly expressed in the target cell types. We also observed that these genes might be highly expressed in non-target cell types, which would affect downstream analysis such as target cell type clustering. Global feature selection in data preprocessing does not remove such local noise because it removes the whole column (cells) or row (genes), potentially leading to substantial information loss. In the denoised gene expression matrix, we observed most of the characterized genes only exhibited high expression levels in their target cell types, whereas their signals became weaker in the non-target cell types. For example, *Bace2* and *Chrdl2* were highly expressed in the endocardial (**Figure 5A** and **B**) and skeletal muscle satellite cells (**Figure 5C** and **D**) in the denoised gene expression matrices, but they were also highly expressed in the cells from non-target cell types in raw gene expression matrices.

### The instance-wise gene selection improves model interpretability

We collected the B cells of brown fat tissues from 3, 18 and 24 months (**Methods**) to investigate if the weight matrix (*W*) could interpret biological observations. For each cell, SUM(*W*) was calculated by aggregating the weights of core genes (**Methods**) as its cell identifier. We found the average values of SUM(*W*) were 15.43, 28.30 and 10.71 for the B cells from 3, 18 and 24 months, suggesting SUM(*W*) could distinguish the cells from different mice age perfectly (**Figure 5E** and **F**).

### Computational performance of IGSimpute

We found a linear relationship between cell number and running time. Increased cell numbers had little effect on the graphics memory usage (**Supplementary Figure 4A** and **B**), suggesting IGSimpute is scalable to imputation on large datasets.

## Discussion

Single-cell sequencing has become an indispensable tool for exploring cell type composition and heterogeneity. To facilitate downstream analysis, it is crucial to ensure that gene expression matrices have accurate values with fewer dropout events and less noise. The existence of dropouts in scRNA-seq data reduces the reliability of various downstream analyses. Imputing these missing gene expression values is therefore an essential task in gene expression matrix preprocessing. Although many imputation methods have been developed, they are commonly influenced by noise and lack interpretability, where all genes have the same weight in dropout imputation.

In this paper, we proposed IGSimpute, an accurate and interpretable deep learning model, to impute the missing values in gene expression profiles derived from scRNA-seq by integrating instance-wise gene selection and gene-gene interaction layers into an autoencoder. The instance-wise gene selection layer efficiently removes noise and highlights the genes that contribute much to the cell embedding. The gene-gene interaction layer explicitly calculates the weights to represent expression dependencies between genes without any distribution assumption. These two components improve the representations of cell-cell and gene-gene correlations, which are the most critical two information sources for dropout imputation. We showed IGSimpute outperformed ten other state-of-the-art tools for imputing dropouts of scRNA-seq derived gene expression data from twelve tissues in the Tabula Muris atlas. Gene expression matrices generated by IGSimpute improved downstream analyses such as cell type clustering and visualization. After post-processing, we found that most marker genes were only highly expressed in the target cell types after imputation. IGSimpute denoised the gene expression matrix by removing outlier genes that were highly expressed in non-target cell types. We found that the gene weights from the instance-wise gene selection layer strongly correlated with the age of B cells.

IGSimpute exhibits linear time complexity with increased cell numbers and imputation can be accelerated with the use of a graphics processing unit (GPU). It is user-friendly and has several additional advantages compared to the methods it was compared to: (i) IGSimpute accepts all types of input gene expression matrices including raw counts, counts per million (CPM), reads per kilobase of exons per million mapped reads (RPKM), fragments per kilobase exons per million mapped fragments (FPKM) and transcripts per million (TPM). This characteristic makes it suitable to be used to re-analyze publicly available scRNA-seq data. (ii) Unlike some methods it was compared to, IGSimpute does not require users to provide a pre-defined number of cell types to guide imputation without introducing extra bias.

IGSimpute has some limitations that should be noted: (i) IGSimpute may not perform well on datasets with few cells (100-1000 cells) due to overfitting in the gene-gene interaction layer. We have however applied regularization techniques, such as global weight decay and dropout layers, to alleviate the influence of low cell numbers. As shown in **Figure 2**, the performance of IGSimpute was acceptable for the heart-and-aorta tissue even though it only included 607 cells; (ii) IGSimpute is not appropriate to be applied to rare cell types (cell proportion < 5%) (**Supplementary Figure 5**). Nevertheless, IGSimpute performed similarly to other methods on rare cell types (**Supplementary Figure 5**). The imputation for rare cell types could be improved on by weighted training (i.e., giving higher weights for the cells from rare cell types). With the development of single-cell ATAC-seq [41–43] and spatial RNA-seq techniques [44–57], it is also possible to extend IGSimpute to these two technologies by considering chromatin accessibility and cell spatial coordinates to improve imputation. scATAC-seq provides additional information on the potential transcription factor binding sites that could be used as prior weights to initialize the gene-gene interaction layer. Moreover, scATAC-seq can help achieve more accurate cell embedding by assuming that cells from the same cell type share similar chromatin accessibilities. The cell coordinates from spatial RNA-seq data will also be helpful in imputation by assuming the cells are from the same cell types if they are close in physical distance. The global gene-gene interaction layer may also be extended to multiple layers for different spatial domains such that each layer can capture gene-gene interaction specific to one spatial domain.

## Data availability

IGSimpute is available at https://github.com/ericcombiolab/IGSimpute. Tabula Muris atlas [33] can be downloaded from https://registry.opendata.aws/tabula-muris/. Tabula Muris Senis atlas [34] can be downloaded from https://registry.opendata.aws/tabula-muris-senis/.

## Key Points

- We present IGSimpute as an accurate and interpretable imputation method with a novel application of instance-wise gene selection for imputing missing scRNA-seq data.
- IGSimpute can impute scRNA-seq data unbiasedly and maintain average expression levels of non-zero entries after imputation.
- We compared IGSimpute with ten other imputation methods using twelve datasets and IGSimpute achieved the lowest average MSE and variance in 9 out of 12 datasets.
- IGSimpute could denoise scRNA-seq data by an interpretable instance-wise gene selection layer and indicate biological characteristics from core genes selected from the layer.

Ke Xu is a Ph.D. student of the Department of Computer Science, Hong Kong Baptist University. His research interests include bioinformatics and computational biology.

ChinWang Cheong is a Ph.D. student of the Department of Computer Science, Hong Kong Baptist University. His research interests include machine learning and health informatics.

Werner Pieter Veldsman is a postdoctoral scholar of Department of Computer Science, Hong Kong Baptist University. His research interests include genomics and bioinformatics.

Aiping Lyu is Chair Professor and Dean of School of Chinese Medicine, Hong Kong Baptist University. His research interests include translational research in Chinese Medicine, drug development and bioinformatics.

William K. Cheung is Professor of Department of Computer Science, Hong Kong Baptist University. His research interests include machine learning and health informatics.

Lu Zhang is Assistant Professor of Department of Computer Science, Hong Kong Baptist University. His research interests include bioinformatics, computational genomics and machine learning.

## Conflict of interest

The authors declare that they have no conflict of interest.

## Author contributions statement

LZ and WC conceived the study; KX and CWC designed IGSimpute; KX and CWC implemented the algorithm, analyzed the results and conducted the experiments; KX, LZ and WV wrote the article; APL and WC reviewed the paper. All authors read and approved the final manuscript.

## Acknowledgment

The authors would like to thank the Shenzhen Science, Technology and Innovation Commission, Research Grants Council of Hong Kong, Hong Kong Baptist University and HKBU Research Committee for their kind support of this project.

## Funding

This research is partially supported by SZVUP Special Fund Project (2021Szvup135), Guangdong-Hong Kong Technology Cooperation Funding Scheme (GHX/133/20SZ), Hong Kong Research Grant Council Early Career Scheme (HKBU 22201419), HKBU Start-up Grant Tier 2 (RC-SGT2/19-20/SCI/007), HKBU IRCMS (No. IRCMS/19-20/D02) and Guangdong Basic and Applied Basic Research Foundation (No. 2021A1515012226).

## Notes

### Competing Interest Statement

The authors have declared no competing interest.

## References

1. Baccin C, Al-Sabah J, Velten L et al. Combined single-cell and spatial transcriptomics reveal the molecular, cellular and spatial bone marrow niche organization. Nat Cell Biol 2020;22(1):38–48.

2. Travaglini KJ, Nabhan AN, Penland L et al. A molecular cell atlas of the human lung from single-cell RNA sequencing. Nature 2020;587(7835):619–25.

3. He B, Chen P, Zambrano S et al. Single-cell RNA sequencing reveals the mesangial identity and species diversity of glomerular cell transcriptomes. Nat Commun 2021;12(1):2141.

4. Morgan D, Tergaonkar V. Unraveling B cell trajectories at single cell resolution. Trends in Immunology 2022;43(3):210–29.

5. Yan M, Hu J, Yuan H et al. Dynamic regulatory networks of T cell trajectory dissect transcriptional control of T cell state transition. Molecular Therapy - Nucleic Acids 2021;26:1115–29.

6. Leader AM, Grout JA, Maier BB et al. Single-cell analysis of human non-small cell lung cancer lesions refines tumor classification and patient stratification. Cancer Cell 2021;39(12):1594–1609.e12.

7. Olalekan S, Xie B, Back R et al. Characterizing the tumor microenvironment of metastatic ovarian cancer by single-cell transcriptomics. Cell Reports 2021;35(8):109165.

8. Ni J, Wang X, Stojanovic A et al. Single-Cell RNA Sequencing of Tumor-Infiltrating NK Cells Reveals that Inhibition of Transcription Factor HIF-1α Unleashes NK Cell Activity. Immunity 2020;52(6):1075–1087.e8.

9. Macosko EZ, Basu A, Satija R et al. Highly Parallel Genome-wide Expression Profiling of Individual Cells Using Nanoliter Droplets. Cell 2015;161(5):1202–14.

10. Svensson V, Vento-Tormo R, Teichmann SA. Exponential scaling of single-cell RNA-seq in the past decade. Nat Protoc 2018;13(4):599–604.

11. See P, Lum J, Chen J et al. A Single-Cell Sequencing Guide for Immunologists. Front. Immunol. 2018;9:2425.

12. Picelli S, Björklund ÅK, Faridani OR et al. Smart-seq2 for sensitive full-length transcriptome profiling in single cells. Nat Methods 2013;10(11):1096–8.

13. Picelli S, Faridani OR, Björklund AK et al. Full-length RNA-seq from single cells using Smart-seq2. Nat Protoc 2014;9(1):171–81.

14. Hashimshony T, Senderovich N, Avital G et al. CEL-Seq2: sensitive highly-multiplexed single-cell RNA-Seq. Genome Biol 2016;17(1):77.

15. Wang X, He Y, Zhang Q et al. Direct Comparative Analyses of 10X Genomics Chromium and Smart-seq2. Genomics, Proteomics & Bioinformatics 2021;19(2):253–66.

16. Andrews TS, Hemberg M. M3Drop: dropout-based feature selection for scRNASeq. Bioinformatics 2019; 35(16):2865–7.

17. Wagner F, Yan Y, Yanai I. K-nearest neighbor smoothing for high-throughput single-cell RNA-Seq data. bioRxiv 2017:217737.

18. Dijk Dv, Sharma R, Nainys J et al. Recovering Gene Interactions from Single-Cell Data Using Data Diffusion. Cell (Cambridge) 2018;174(3):716–729.e27.

19. Huang M, Wang J, Torre E et al. SAVER: gene expression recovery for single-cell RNA sequencing. Nature methods 2018;15(7):539–42.

20. Li WV, Li JJ. An accurate and robust imputation method scImpute for single-cell RNA-seq data. Nature communications 2018;9(1):997.

21. Hu Y, Li B, Zhang W et al. WEDGE: imputation of gene expression values from single-cell RNA-seq datasets using biased matrix decomposition. Brief Bioinform 2021;22(5):

22. Eraslan G, Simon LM, Mircea M et al. Single-cell RNA-seq denoising using a deep count autoencoder. Nat Commun 2019;10(1):390.

23. Arisdakessian C, Poirion O, Yunits B et al. DeepImpute: an accurate, fast, and scalable deep neural network method to impute single-cell RNA-seq data. Genome biology 2019;20(1):211.

24. Amodio M, van Dijk D, Srinivasan K et al. Exploring single-cell data with deep multitasking neural networks. Nat Methods 2019;16(11):1139–45.

25. Wang J, Ma A, Chang Y et al. scGNN is a novel graph neural network framework for single-cell RNA-Seq analyses. Nat Commun 2021;12(1):1882.

26. Lopez R, Regier J, Cole MB et al. Deep generative modeling for single-cell transcriptomics. Nat Methods 2018;15(12):1053–8.

27. Wolf FA, Angerer P, Theis FJ. SCANPY: large-scale single-cell gene expression data analysis. Genome Biol 2018;19(1):15.

28. Satija R, Farrell JA, Gennert D et al. Spatial reconstruction of single-cell gene expression data. Nat Biotechnol 2015;33(5):495–502.

29. Zheng GXY, Terry JM, Belgrader P et al. Massively parallel digital transcriptional profiling of single cells. Nat Commun 2017;8(1):14049.

30. Stuart T, Butler A, Hoffman P et al. Comprehensive Integration of Single-Cell Data. Cell 2019;177(7):1888–1902.e21.

31. Yoon J, Jordon J, van der Schaar M. INVASE: Instance-wise Variable Selection using Neural Networks. International Conference on Learning Representations 2018.

32. Masoomi A, Wu C, Zhao T et al. Instance-wise Feature Grouping. Advances in Neural Information Processing Systems 2020;33:13374–86.

33. Single-cell transcriptomics of 20 mouse organs creates a Tabula Muris. Nature 2018;562(7727):367–72.

34. A single-cell transcriptomic atlas characterizes ageing tissues in the mouse. Nature 2020;583(7817):590–5.

35. Jiang L, Schlesinger F, Davis CA et al. Synthetic spike-in standards for RNA-seq experiments. Genome Res. 2011;21(9):1543–51.

36. Abadi M, Agarwal A, Barham P et al. TensorFlow: Large-Scale Machine Learning on Heterogeneous Distributed Systems, 2016.

37. Loshchilov I, Hutter F. Decoupled Weight Decay Regularization, 2017.

38. Kiselev VY, Kirschner K, Schaub MT et al. SC3: consensus clustering of single-cell RNA-seq data. Nat Methods 2017;14(5):483–6.

39. Hubert L, Arabie P. Comparing partitions. Journal of Classification 1985;2(1):193–218.

40. Becht E, McInnes L, Healy J et al. Dimensionality reduction for visualizing single-cell data using UMAP. Nat Biotechnol 2018;37(1):38–44.

41. Buenrostro JD, Wu B, Litzenburger UM et al. Single-cell chromatin accessibility reveals principles of regulatory variation. Nature 2015;523(7561):486–90.

42. Cusanovich DA, Daza R, Adey A et al. Multiplex single-cell profiling of chromatin accessibility by combinatorial cellular indexing. Science 2015;348(6237):910–4.

43. Corces MR, Trevino AE, Hamilton EG et al. An improved ATAC-seq protocol reduces background and enables interrogation of frozen tissues. Nat Methods 2017;14(10):959–62.

44. Chen KH, Boettiger AN, Moffitt JR et al. Spatially resolved, highly multiplexed RNA profiling in single cells. Science 2015;348(6233):aaa6090.

45. Ståhl PL, Salmén F, Vickovic S et al. Visualization and analysis of gene expression in tissue sections by spatial transcriptomics. Science 2016;353(6294):78–82.

46. Orjalo AV, Johansson HE. Stellaris® RNA Fluorescence In Situ Hybridization for the Simultaneous Detection of Immature and Mature Long Noncoding RNAs in Adherent Cells. In: Feng Y, Zhang L (eds). Long Non-Coding RNAs: Methods and Protocols. New York, NY: Springer New York, 2016, 119–34.

47. Moffitt JR, Hao J, Wang G et al. High-throughput single-cell gene-expression profiling with multiplexed error-robust fluorescence in situ hybridization. PNAS 2016;113(39):11046–51.

48. Chen J, Suo S, Tam PP et al. Spatial transcriptomic analysis of cryosectioned tissue samples with Geo-seq. Nat Protoc 2017;12(3):566–80.

49. Shah S, Takei Y, Zhou W et al. Dynamics and Spatial Genomics of the Nascent Transcriptome by Intron seqFISH. Cell 2018;174(2):363–376.e16.

50. Nichterwitz S, Benitez JA, Hoogstraaten R et al. LCM-Seq: A Method for Spatial Transcriptomic Profiling Using Laser Capture Microdissection Coupled with PolyA-Based RNA Sequencing. In: Gaspar I (ed). RNA Detection: Methods and Protocols. New York, NY: Springer New York, 2018, 95–110.

51. Codeluppi S, Borm LE, Zeisel A et al. Spatial organization of the somatosensory cortex revealed by osmFISH. Nat Methods 2018;15(11):932–5.

52. Xia C, Fan J, Emanuel G et al. Spatial transcriptome profiling by MERFISH reveals subcellular RNA compartmentalization and cell cycle-dependent gene expression. PNAS 2019;116(39):19490–9.

53. Vickovic S, Eraslan G, Salmén F et al. High-definition spatial transcriptomics for in situ tissue profiling. Nat Methods 2019;16(10):987–90.

54. Rodriques SG, Stickels RR, Goeva A et al. Slide-seq: A scalable technology for measuring genome-wide expression at high spatial resolution. Science 2019;363(6434):1463–7.

55. Eng C-HL, Lawson M, Zhu Q et al. Transcriptome-scale super-resolved imaging in tissues by RNA seqFISH+. Nature 2019;568(7751):235–9.

56. Stickels RR, Murray E, Kumar P et al. Highly sensitive spatial transcriptomics at near-cellular resolution with Slide-seqV2. Nat Biotechnol 2021;39(3):313–9.

57. Xia K, Sun H-X, Li J et al. The single-cell stereo-seq reveals region-specific cell subtypes and transcriptome profiling in Arabidopsis leaves. Developmental Cell 2022.

