## Supplementary Materials for "Accurate and interpretable gene expression imputation on scRNA-seq data using IGSimpute"

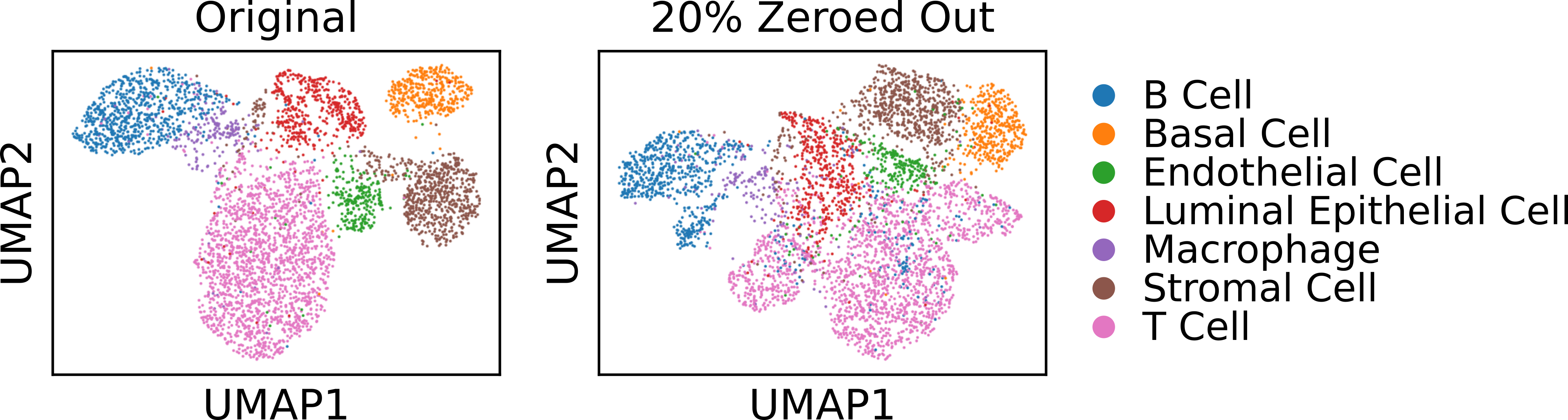

**Supplementary Figure 1.** UMAP was performed to visualize mammary gland gene expression before and after zeroing out 20% of non-zero entries. Colors represent the respective cell types. Zeroing out noticeably increased cluster mixtures in comparison to the original profiles. (**Left**) The UMAP plot of the original profiles has discernable delineated boundaries. (**Right**) The UMAP plot of the 20% zeroed-out profiles. Inter-cluster distances become shorter and some B cells are mixed with T cells. The original B cell cluster is split into two subclusters and the original T cells cluster is split into three subclusters.

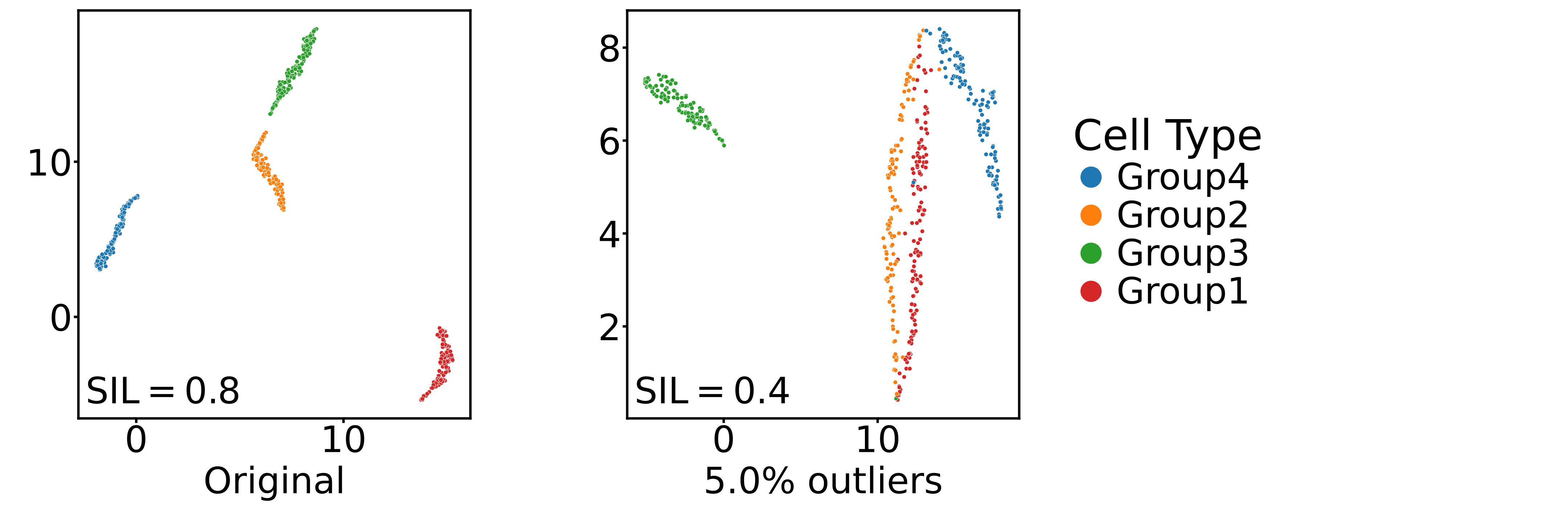

**Supplementary Figure 2.** Outliers led to inaccurate cell-cell distance estimation and resulted in mixed clusters. (**Left**) The UMAP plot of 1,000 simulated cells without outliers. (**Right**) The UMAP plot of 1,000 simulated cells with 5% outliers. SIL, silhouette score.

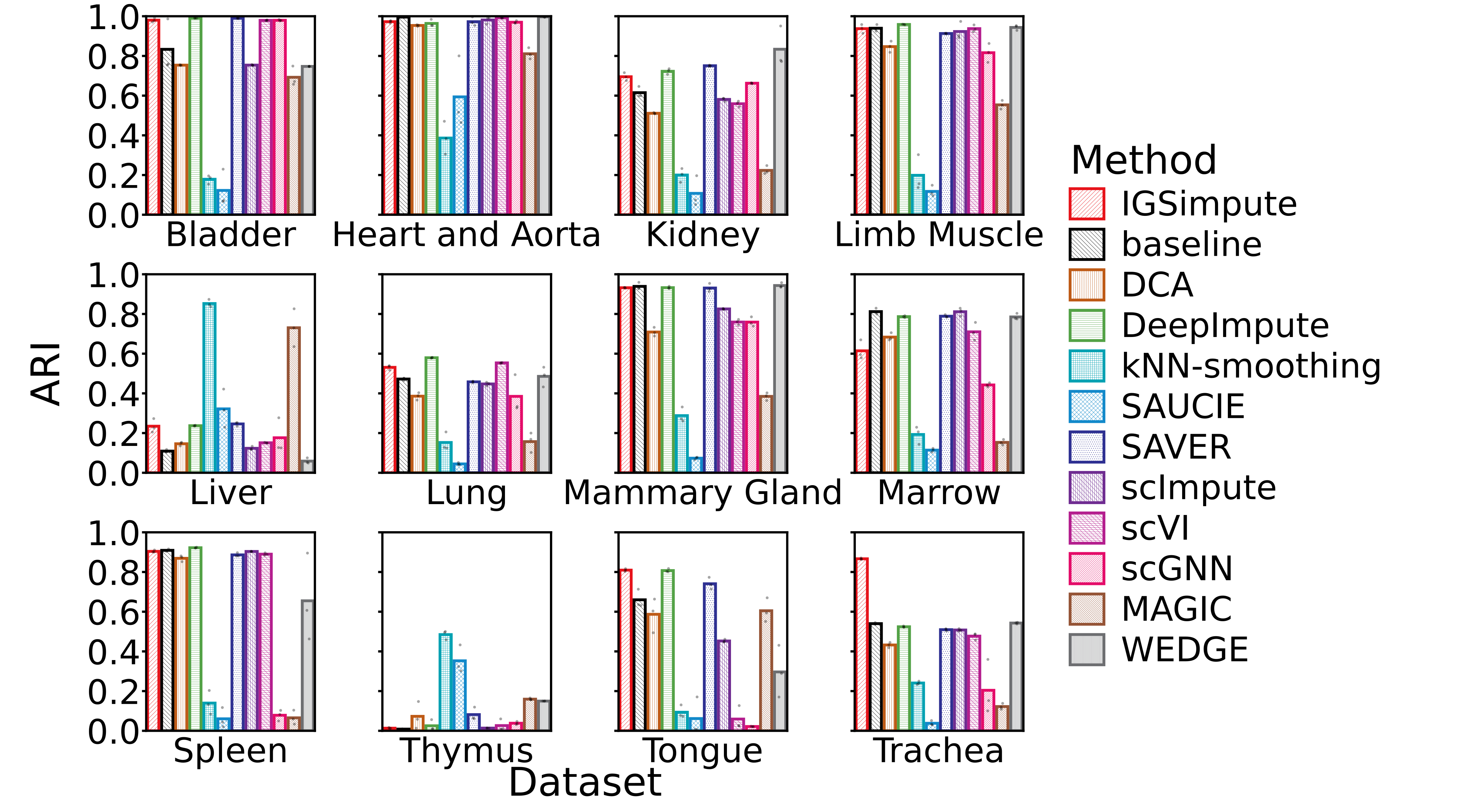

**Supplementary Figure 3.** The clustering results of all imputed profiles for the 12 tissues in the Tabula Muris atlas. The bar plots illustrate replication results for all tissue datasets and methods. Each bar represents the median value for the clustering results. For most of the methods and datasets, the same parameters were used when performing clustering by SC3. ARI, adjusted rand index.

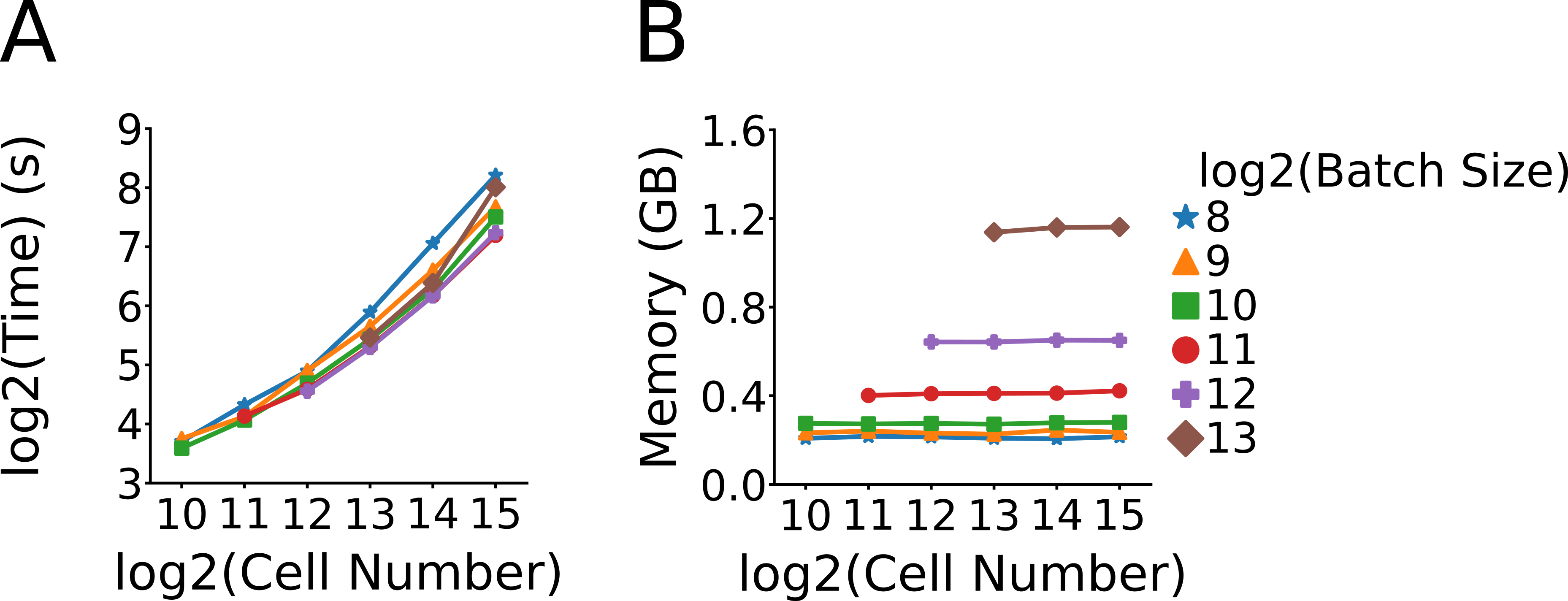

**Supplementary Figure 4.** IGSimpute is scalable to larger datasets. (**A**) Running time increases linearly with cell number. (**B**) Graphics memory usage increases linearly with cell number.

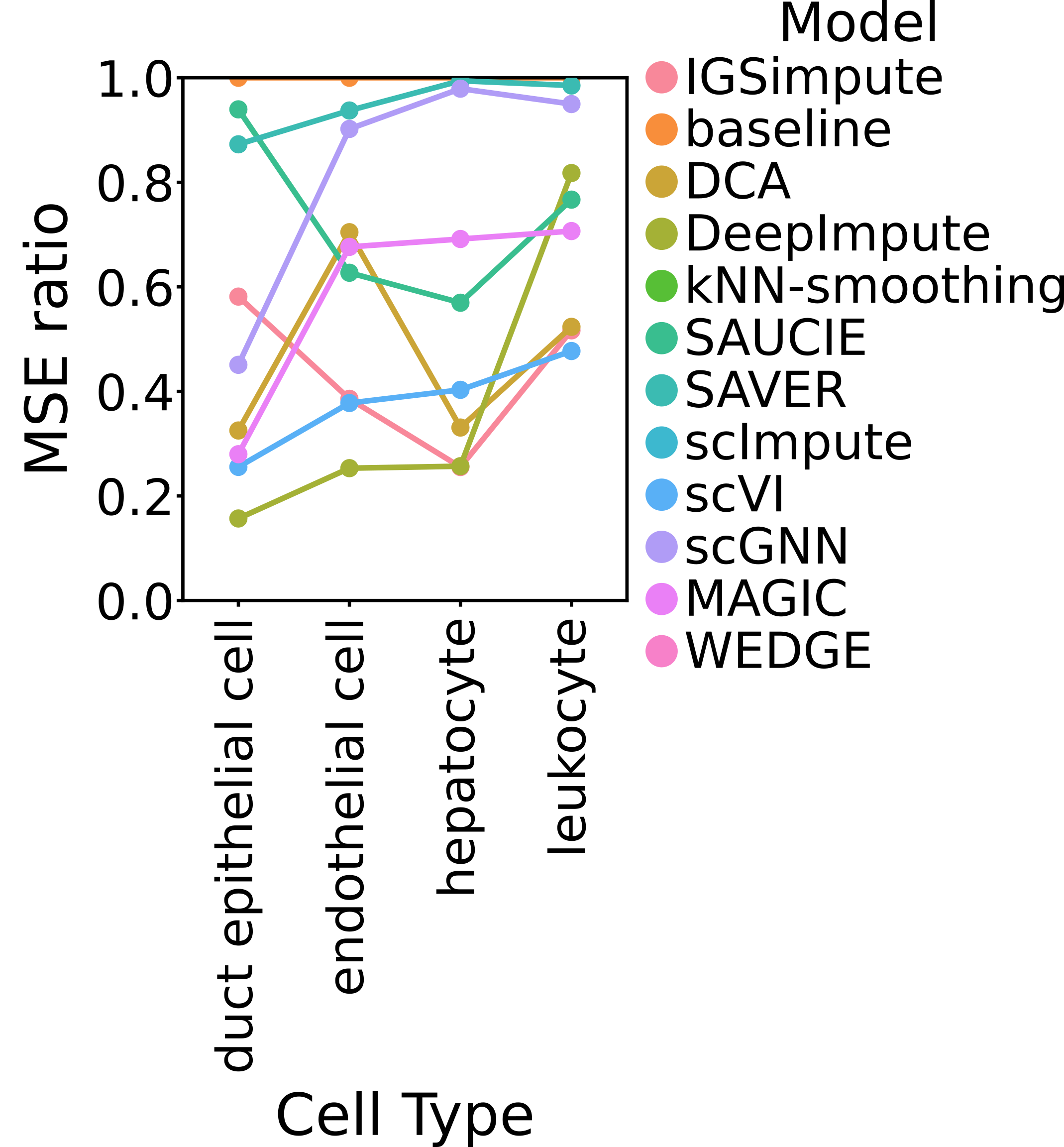

**Supplementary Figure 5.** Imputation performance grouped by cell type. IGSimpute underperformed on rare cell types compared to common cell types. Duct epithelial cells, endothelial cells or leukocyte cells accounted for less than 2% of all cells. MSE ratio, (MSE of current method) / (MSE of baseline).

**Supplementary Table 1.** Dataset information

| **Dataset name** | **Number of cells** | **Number of genes** | **Number of cell groups** |
| --- | --- | --- | --- |
| Bladder | 2500 | 23433 | 4 |
| Heart_and_aorta | 607 | 23433 | 5 |
| Kidney | 2781 | 23433 | 8 |
| Limb_muscle | 3909 | 23433 | 6 |
| Liver | 1845 | 23433 | 4 |
| Lung | 5404 | 23433 | 13 |
| Mammary_gland | 4481 | 23433 | 7 |
| Marrow | 3652 | 23433 | 14 |
| Spleen | 9552 | 23433 | 5 |
| Thymus | 1429 | 23433 | 3 |
| Tongue | 7538 | 23433 | 3 |
| Trachea | 11269 | 23433 | 5 |
| Brown adipose tissue (BAT) | 7896 | 7824 | 1 |

**Supplementary Note 1.** Dropout simulation and outlier simulation.

For dropout simulation, we used Splatter to simulate two groups of cells with each group consisting of 500 cells with 2,000 genes. Entries with gene expression values equal to six would have 50% dropout rates, resulting in approximately 70% dropout rates for all entries. The outlier probability was set to 5%. For outlier simulation, we used Splatter to simulate four groups of cells with each group consisting of 250 cells with 2,000 genes. The outlier probabilities were set to 0 and 5%, respectively. The outlier factor scale was adjusted to 1.

**Supplementary Note 2.** Dataset information.

*Droplet-based Tabula Muris Datasets*

Tabula Muris is an atlas of mouse single-cell transcriptomics and contains gene count tables of 12 tissues constructed using the 10x Genomics protocol. The gene count table in h5ad format and corresponding cell type labels in CSV format were downloaded from Amazon Web Services according to the guidance of https://github.com/czbiohub/tabula-muris/blob/master/tabula-muris-on-aws.md. We treated the "tissue" column in the CSV file as the tissue labels for dividing the atlas into different datasets containing one pure tissue each. We extracted "cell_ontology_class" as the ground truth labels in the clustering analysis.

*Brown Adipose Tissue in the Facs-based Tabula Muris Senis Dataset*

Tabula Muris Senis is an atlas of single-cell transcriptomics that depicts the life span of mice and contains gene count tables of 18 tissues constructed using the 10x Genomics protocol. The gene count table together with age and cell types annotation in h5ad format was downloaded from https://figshare.com/projects/Tabula_Muris_Senis/64982. We treated the "tissue" column in the obs attribute of the annData object loaded from h5ad by Scanpy as the tissue labels for extracting the brown adipose tissue (BAT) from the atlas. We subsequently filtered B cell data from the BAT data using terms in the "cell_ontology_class" column. The "age" column as the ground truth in all experiments was never exposed to IGSimpute and was only used in representing and analyzing results.

**Supplementary Note 3.** Detailed structure of IGSimpute.

IGSimpute has integrated autoencoders with a gene selection autoencoder for embedding and a gene-gene interaction layer for decoding. The encoder accepts normalized scRNA-seq data via its input layer (*Input*). This is followed by a Gaussian noise layer (*Gauss*) which applies a 1.5 standard deviation for learning denoised representations. A five-layered selection autoencoder is connected with *Gauss* for selecting informative genes. The first three layers of the selection autoencoder (*Selection_encoder, Selection_embedding, Selection_decoder*) have output dimensions of 400, 200, 400 respectively and employ ReLU as the activation function. The fourth layer (*Selection_selector*) acts as the gene selection layer. Its output dimension is equal to the number of genes, and it employs the sigmoid function as the activation function. The fifth layer (*Selection_dropout*) is a dropout layer with a 20% dropout rate for learning robust representations. The outputs of *Gauss* and *Selection_dropout* are multiplied and passed through a fully-connected layer (*Encoder_FC*) with an output dimension of 400. *Encoder_FC* does not employ an activation function. The output of *Encoder_FC* can be seen as the output of the final embedding layer, while the following layers can be seen as part of the decoder. A fully-connected layer, *Decoder_FC*, is connected with *Encoder_FC.* It has an output dimension equal to the number of genes, and it employs a Softplus activation function for constructing the initial imputed expression matrix $\tilde{X}$. Three learnable tensors and a non-zero mask, matrix$M$, are introduced for finetuning the imputation results. The first learnable tensor is a gene-gene interaction matrix$G$, where the diagonal entries are coerced to zero in order to negate trivial interactions such as genes interacting with themselves. Its row length and column length are the same as the number of genes. The second learnable tensor is a bias vector $b$ with a length equal to the number of genes. It employs a ReLU activation function for quantifying the expression values that are not interpreted by the interaction matrix. The third learnable tensor is a weight vector$p$ with the same length as the second learnable tensor but with a sigmoid activation function for controlling the explained ratio of the imputed values by the gene-gene interactions or the bias vector. The imputed expression matrix from the network is as follows:

$$\hat{X}=p\odot\left( \left( \tilde{X}\odot\left( 1-M \right)+X\odot M \right)\cdot G \right)+(1-p)\odot b$$

where $X$ is the count matrix.

**Supplementary Note 4.** Imputation software settings.

*IGSimpute*: Normalization was done as illustrated in **Method**. $\lambda_{gg}$, $\lambda_{rec}$ and $\lambda_{gs}$ were set to 1, 0.1 and 0.01 respectively.

*DCA*: Library size normalization, gene-wise normalization and log transformation were performed in order.

*DeepImpute*: Internal normalization during the fitting step and prediction step was allowed.

*kNN-smoothing*: K was set to 10 to avoid over-smoothing on rare cell types. The D parameter was kept at its default value of 10.

*MAGIC*: The number of principal components was kept at its default value of 100.

*SAUCIE*: The input data for training and testing were identical to that used in the SAUCIE tutorial.

*SAVER*: SAVER was used in single-threaded mode since the multithread mode was error-prone. The size factor also had to be set to one in order to avoid normalized profiles instead of count profiles being returned.

*scGNN*: We carried out a pre-processing step using a left-truncated mixture Gaussian (LTMG) distribution as recommended by the scGNN tutorial and we run scGNN in non-sparse mode since comma-separated values (CSV) format was used. The quick mode was not adopted for better performance though it could reduce running time significantly. The ground truth cluster number was given for internal clustering in order to obtain better clustering results. During the eighth imputation replicate on tongue tissue data from the Tabula Muris atlas, the quick mode had to be turned on to avoid an internal bug that produced an empty intermediate variable and prevented scGNN from running. LTMG and the quick mode were also turned off during the seventh imputation replicate on heart and aorta tissue data for the same reason.

*scImpute*: Cell type labels were not provided but ground truth cluster numbers were provided.

*scVI*: The standalone scVI package was adopted because the scVI package integrated into the scvi-tools package only produced normalized profiles. The number of epochs was set to 400 and the learning rate was set to 0.001.

*WEDGE*: The MATLAB version of WEDGE was used with multithread acceleration. Library size normalization and log transformation were carried out internally.
